# Impact of female mate copying on male morph dynamics

**DOI:** 10.1101/2022.09.13.507757

**Authors:** Srishti Patil, Sabine Nöbel, Chaitanya S. Gokhale

## Abstract

Mate copying (MC), a type of non-independent mate choice, is a behaviour observed in many vertebrate and few invertebrate species. It occurs when an individual’s sexual preference gets socially inclined toward those of its conspecifics. Theoretical models and experimental studies of MC have been limited to choice between two options (or morphs). In this study we model the evolution of morphology in a population under varying extents of mate copying. Multiple morphs and multiple observations are considered and a generalisable model is presented. We quantify the level of copying needed to achieve pseudo-stable equilibria in the presence of multiple morphs. Moving closer to realistic scenarios we support our theoretical development with simulations and discuss relevance for empirical model systems.

## Introduction

A basic assumption of classical theories of mate choice is that an individual’s identification and selection of a high-quality mate is independent of other conspecifics. However, the signals and cues targeted toward the chosen mate may provide public information to other individuals. A growing body of evidence shows that this social information may affect mate choice, resulting in what is called non-independent mate choice [1, 2, 3, 4]. The net effect is that individuals may use personal information, social information or both to facilitate mate choice.

Mate copying (MC), a subset of non-independent mate choice, is a behaviour in which an individual copies the mate choice of conspecifics when choosing a mate. Mate copying has been experimentally demonstrated in a large number of species of vertebrates including birds [5, 6, 7], mammals [8, 9], and fish [10, 11]. The first invertebrate in which it was observed is the common fruit fly, *Drosophila melanogaster*. Mery and colleagues demonstrated MC in *D. melanogaster* using artificially generated male phenotypes by dusting them with pink and green powders [2]. In the last decade, many experiments and empirical studies have been carried out to test the effect of various conditions on the mate-copying behaviour of fruit flies, like atmospheric pressure [12] and sex ratio [13].

Studying MC is important because it affects sexual selection and can play a role in significant processes like speciation and hybridisation [14, 15, 16]. The evolutionary consequences of this behaviour have been tested empirically and modelled in the case with two male morphs, which means that individual females have one choice between two males. In the case of [2], the choice was between green males and pink males, and this choice was affected by social learning. When the two choices have different fitness or “qualities”, we expect to see interesting effects on the population dynamics. In *D. melanogaster*, it has been shown that MC could lead to the preference of a male with lower quality over the one with higher quality [17]. In this study, we ask if MC can maintain polymorphism in populations in the presence of multiple morphs.

It is known that MC helps the spread of a fitter novel trait [18], which suggests an explanation as to why MC would evolve and how it could be adaptive. Building on that, a mechanism for the evolution of MC (given by [19]), shows how MC is adaptive even when the copying allele is itself mildly deleterious. On the other hand, game theory suggests that social information use akin to MC can lead to maladaptive decisions ([20], see also [21]). Therefore, MC under different contexts leads to different costs and benefits and consequently, differences in the proportion of copying individuals, which can also be interpreted as the probability of copying. We use the probability of copying as a parameter to study the effects of varying degrees of female MC on the male population. We assume the copying individuals to be females, although MC has been demonstrated in males of some species [22].

Previous theoretical studies on MC, among other things, have modelled the evolution of mate copying under various assumptions about development of preference, collection of information and interactions between individuals [23, 24, 25, 26]. It requires a careful analysis of costs and benefits of copying versus choosing based on personal information [27], which in turn depend on many ecological factors like sex ratio, population density, spatial distribution and age distribution. For instance, MC in *D. melanogaster* females correlates with atmospheric pressure in certain situations [12]. While our theory development is not species specific, we take *D. melanogaster* as the prime example to develop outwards from.

Our model builds and expands on the previous theoretical models of MC [28, 15, 24, 19, 23]. Firstly, the nature and manner of collection of information become important because they affect the production of the behaviour. We consider two classes of information - personal and public (social). Personal information is independent of choices made by other individuals and leads to inherent preferences. Social or public information is learnt and depends on the behaviour of conspecifics leading to cultural preferences. This differentiation is necessary because MC can cause directional or frequency-dependent selection, depending on how social information influences females [28]. Therefore to get a complete understanding, we need to analyse how social information interacts with ecological and life-history variables [3]. If not accounted for, these variables can lead to seemingly contrasting results. An example is the effect of the number of observations on the copying behaviour. A simulation study by [16] predicts that more observations and copying in a conformist manner result in a greater stability of a tradition, which could be interpreted as a higher degree of cultural influence. However, the mathematical model proposed by Agrawal [28] predicts that more observations imply less copying and hence less cultural influence. The difference is in the assumption of how information is obtained and used by individuals.

Second, we consider the nature of matings. This involves sexual conflict [29, 30] and male-male competition [31]. We take into account how these mating complexities affect the outcome of mate choice, which depends on which type of information (personal or public) is dominant. The ceiling effect - diminished effect of information in favour of the inherently preferred morph - clearly demonstrates a kind of asymmetry in the way inherent and cultural preferences are formed [17].

A step towards understanding how public information is processed by individuals was the link between MC and conformism in *D. melanogaster* [16]. Conformism is the exaggerated tendency to copy the majority [32] and mate copying, at least in *D. melanogaster*, is a conformist behaviour [16]. This result alone makes it possible to apply many insights from studies on conformist behaviour to a biological system like mating in *D. melanogaster*. For instance, different ways to evaluate “majority” may lead to varying outcomes in conformist decision-making [33]. A wide range of results about conformist biases in cultural transmissions and how they interact with other evolutionary forces like natural selection [34], can be applied.

Experimentation on mating systems with MC presents the difficulty in observing and disentangling learning and evolutionary timescales. Theoretical studies are crucial at that juncture and help us make predictions about the evolutionary fate of the population. Our study contributes to this respect by proposing a model that describes the evolutionary trajectories of populations with multiple morphs under the effect of mate copying. Our study can help obtain conditions under which a conformist behaviour like MC leads to the desired evolutionary outcomes. Theoretical development of MC including more than two morphs has not been explored yet. The following sections describe the model and its results for two separate cases and show how MC can lead to polymorphism in a population.

## Model & Results

Mate preferences can drastically affect the dynamics of a population [20, 15]. Our model assumes two kinds of mate preferences - an inherent evolutionarily determined preference depending on the quality of a male morph, and mate copying. Preference for a male morph would increase its frequency in the population. Similarly, preference against a morph would reduce its proportion in the population. We represent this modification of the quality and, subsequently, the growth rate of male morph *i* with the “preference factor”, denoted by *P*_*i*_. The row vector P contains all the preference factors.

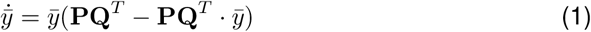

This describes a system of equations for the dynamics of the male population (Eq (1)), where 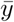 is the frequency vector and Q = [*q*_1_, *q*_2_, …, *q*_*m*_] for *m* male morphs. We use the term “quality” to refer to the classical concept of fitness, i.e. the average number of offspring. In the equation, we modify the quality by multiplying it to the preference factor (*P*_*i*_*q*_*i*_), which we refer to as male fitness. Therefore, male quality is independent of female preference, but male fitness (or modified quality) is not. This reformulation helps us take the female preferences into account. The preference factors are given by,

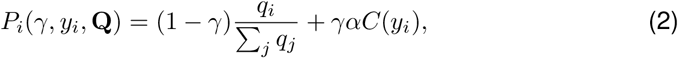

where *γ* is the probability of copying. We introduce the factor of asymmetry *α* because the numerical values and weights given to inherent and cultural preferences may not be comparable. In equation (2), the first term represents inherent preference, which is proportional to the relative quality and the second term accounts for mate copying.

In a population of females that display mate copying behaviour, we assume that every individual will choose to copy with a probability *γ*. When copying occurs, the female will switch its preference to a male morph observed to be mating with other females. The probability of switching to male morph *i* is then given by the “copying function” (Eq. (3)). If the female chooses not to copy, mating happens based on inherent preference. This form of the preference factor (Eq. (2)) assumes an infinite, well-mixed population.

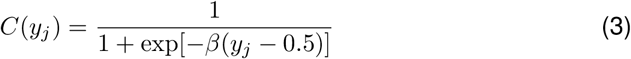

### Training Period

The copying function *C* is solely dependent on the frequency of matings involving male morph *j*, which in our case can be approximated by the frequency *y*_*j*_ of the morph in the entire male population. How do the females obtain and learn this information? We assume an initial training period in each generation which is a fraction of the total number of matings in that generation (*n* matings, a fraction *u* of the total number of matings) [28]. There is no mate copying in the training period (*γ* = 0), and all females observe all the matings. Since the training period occurs in every generation, an additional term is added to equation (1) with *γ* = 0. Thus in every generation, females use the information learnt from the initial *n* matings in the training period for decision-making in the copying period (Figure 1). The following equation (where P^0^ is the vector of preference factors with *γ* = 0.) includes the training period.

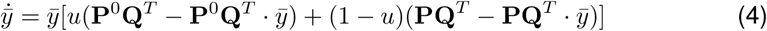

**Figure 1:**
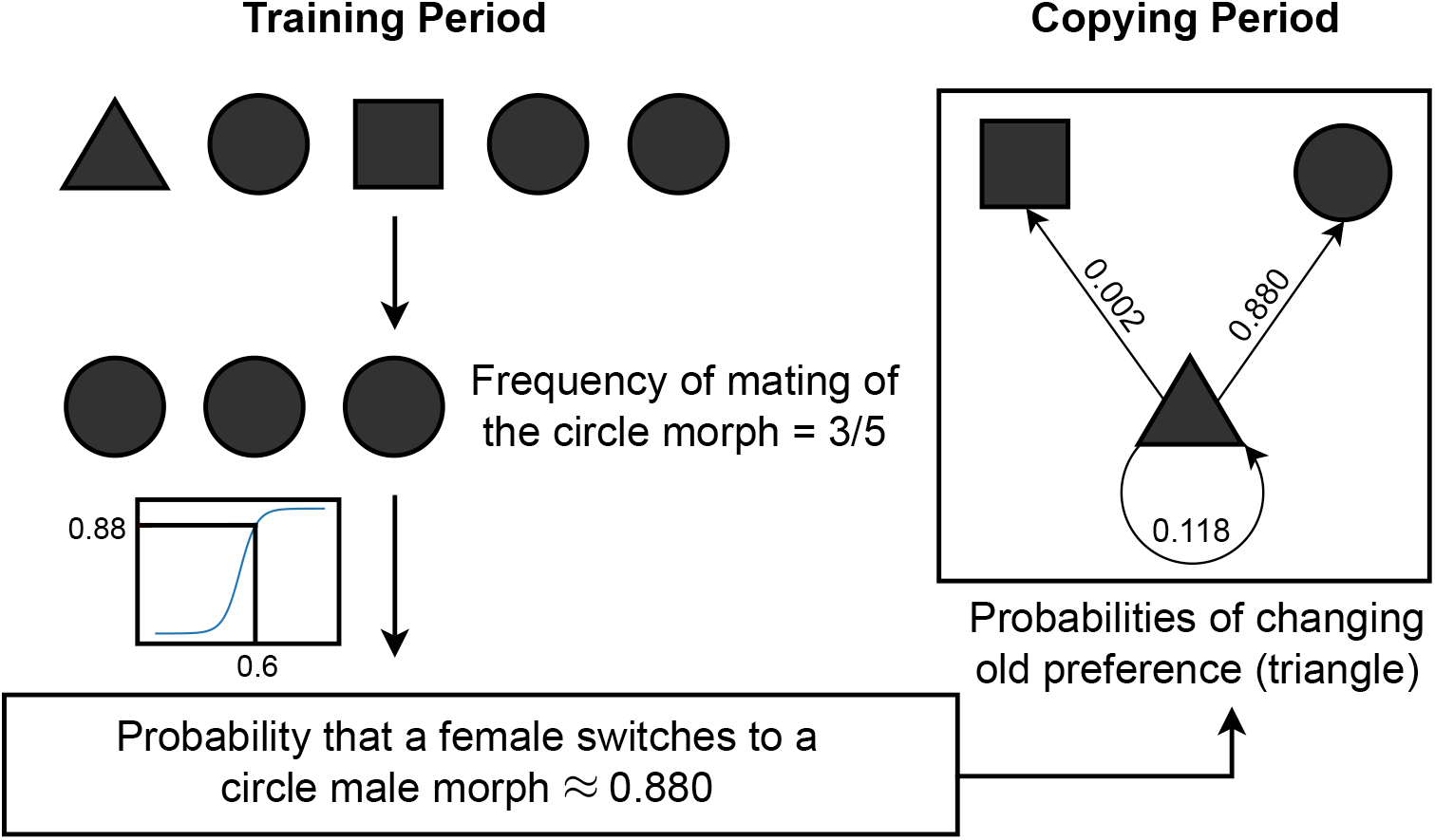
Illustration of the training and copying periods for a system with female mate copying. There are three male morphs (denoted by triangles, circles and squares) and five matings (or observations) in the training period. The probability of switching preference to a circle morph in the copying period is given by the value of the function *C* at 3*/*5. Similarly, the probabilities of switching from the triangle morph to the other two morphs are calculated. These probabilities come from the values of the copying function *C*(*f*_*i*_) where *f*_*i*_ is the frequency of the *i*^th^ morph in the training period. Hence, information collected during the training period is used to decide the switching probabilities in the copying period.

### Conformism

Mate copying can be a conformist behaviour [16], i.e. there can be an exaggerated tendency for individuals to copy the majority [32]. In practice, it implies that *C*(*y*) *>* with the more straightforward case*y* when *y >* 0.5 and *C*(*y*) *< y* when *y <* 0.5. In equation (3), *β* can be interpreted as the extent of conformism. For negative values of *β*, we get anti-conformism.

It is essential to recognise the difference between *β* and *γ* interpretations. The extent of conformism *β*, is a parameter to specify the form of the conformism function. It is fixed at a constant value throughout the analysis allowing us to observe conformism. Changing *β* amounts to changing the difference between *y* and *C*(*y*). Note that *β* = 0 does *not* correspond to random choice. On the other hand, *γ* represents the extent of copying, specifying the amount of weight given to inherent preference over of cultural preference. When female preference is completely determined by quality (or inherent preference) and is not influenced by the frequency of the male morph, then *γ* = 0.

### Two Morphs

We begin with the more straightforward case of two morphs to exemplify the model dynamics. We solve equation (4) for two male morphs with different quality values. Let *y*_*t*_ be the frequency of the lower-quality morph at time *t*. For any value of *γ*, there are at least 2 fixed points, at *y* = 0 and at *y* = 1. A third fixed point appears for a particular range of parameter values (Figure 2).

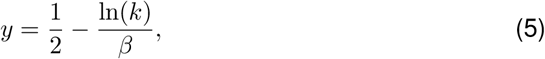

where *k <* 1 is a function of the parameters *γ, α*, Q and *u* (see Section S.1 for derivation). From equation (5), we see that any lower quality morph with an initial frequency below 0.5 cannot reach fixation in the male population. To verify the results, we ran simulations with specific values of parameters for different initial conditions (Figure 2c, see Table 2 for parameter values).

**Figure 2:**
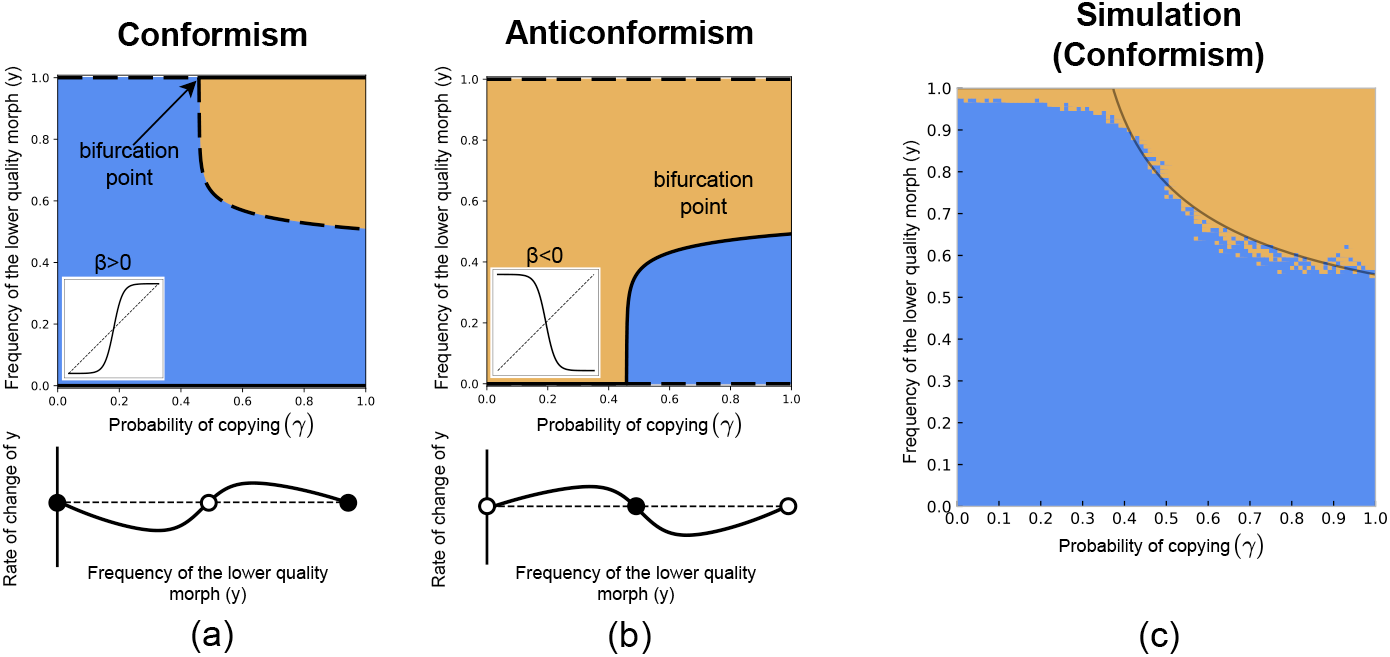
**(a)**,**(b)** Bifurcation diagrams for conformist and anti-conformist mate copying (Q = [2.0, 2.3], *α* = 0.2, *u* = 0.01, *β* = ±20). Dashed lines are unstable fixed points and solid lines are stable fixed points. The frequency of the lower-quality morph is *y*. The coloured regions mark the basins of attraction for the fixed points *y* = 0(blue) and *y* = 1(orange). Thus, any initial point lying in the orange region will have the lower quality morph going to fixation. The probability of switching to the lower-quality male morph is given by *C*(*y*) (insets), which is a function of the extent of conformism, *β*. Positive values display conformist behaviour and negative values indicate anti-conformism. The dynamics of the low-quality morph frequency in a two-morph population give rise to stable(unstable) fixed points at the extremes and unstable(stable) fixed point in the middle for *β >* 0(*β <* 0). **(c)** Results of the simulations run for the two-morph system (described in Section S.3) with different initial conditions. The blue(orange) points are initial conditions from where the higher(lower) quality morph goes to fixation. Simulation parameters are listed in Table 2. The grey line separating the two basins of attraction is obtained by setting *β* = 4 while the simulation used *β* = 20 to compute the probability of switching. This discrepancy is addressed in Section Search Parameter and Effective Conformism

**Table 1:** List of model parameters

| Parameter | Description |
| --- | --- |
| $y_i$ | Frequency of the $i^{\text{th}}$ male morph |
| $q_i$ | Quality of the $i^{\text{th}}$ male morph |
| $\alpha$ | Factor of asymmetry |
| $\beta$ | Extent of conformism |
| $\gamma$ | Probability of copying |
| $m$ | Number of male morphs |
| $u$ | Fraction of matings in the training period |

**Table 2:**
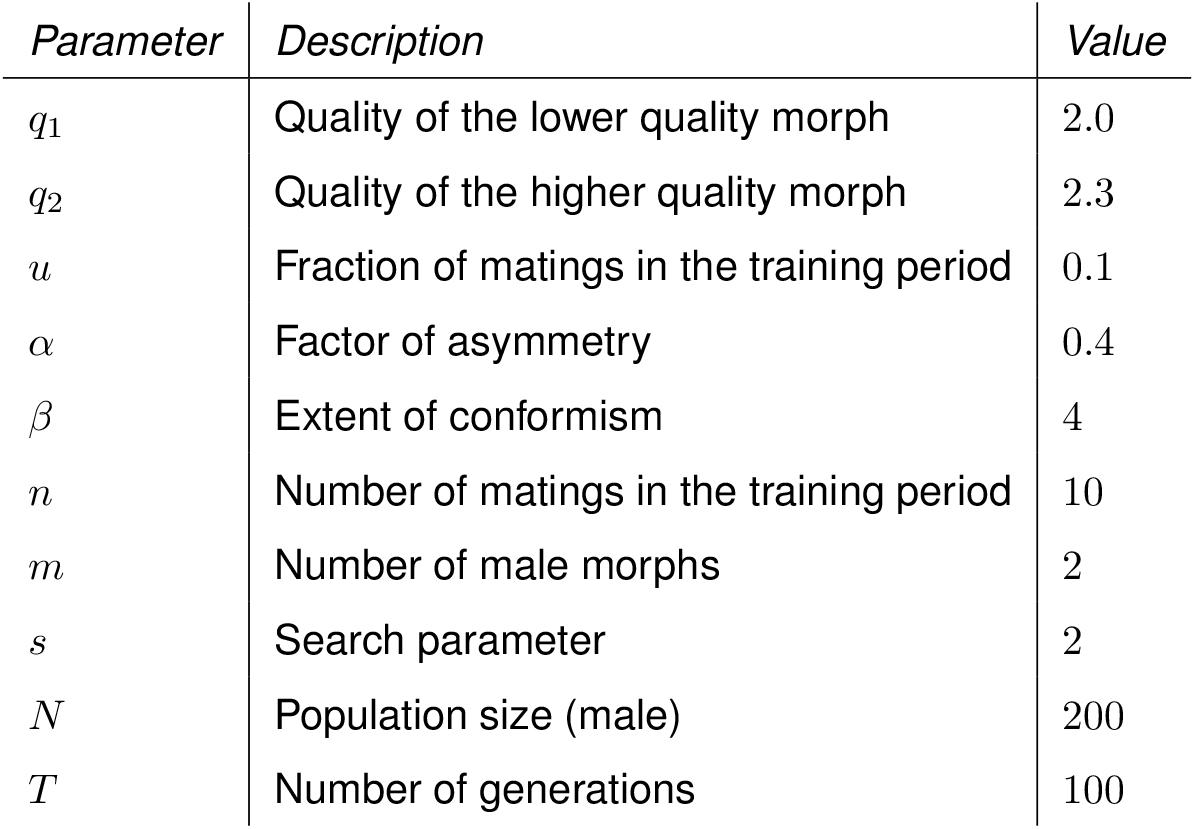
List of simulation parameters with values

There are two basins of attraction in the *γ* − *y* phase space. After the point of bifurcation (when the third fixed point appears), the lower quality morph can fix in the population. The time taken to reach fixation, obtained numerically, is relatively low in these regions but increases as we approach the separatrix defined by the set of unstable fixed points (compare separatrix position in Figure 2a and Figure 3a). Fixation time can be correlated to the absolute difference between the modified qualities (i.e. PQ^*T*^) of the two morphs (Figure 3b). When this quality difference is high, there is a strong preference for one of the morphs due to mate copying or inherent preference. This results in a significant difference in growth rates of the two male subpopulations and hence a faster approach to fixation. When the quality difference is low, the growth rates are similar, with more competition among the two male morph subpopulations. Looking at the basins of attraction and the absolute difference between modified qualities, we know that there must be a set of points where this difference goes to zero, separating the two regions. This hints at the possibility of speciation due to mate copying [14] under specific conditions.

**Figure 3:**
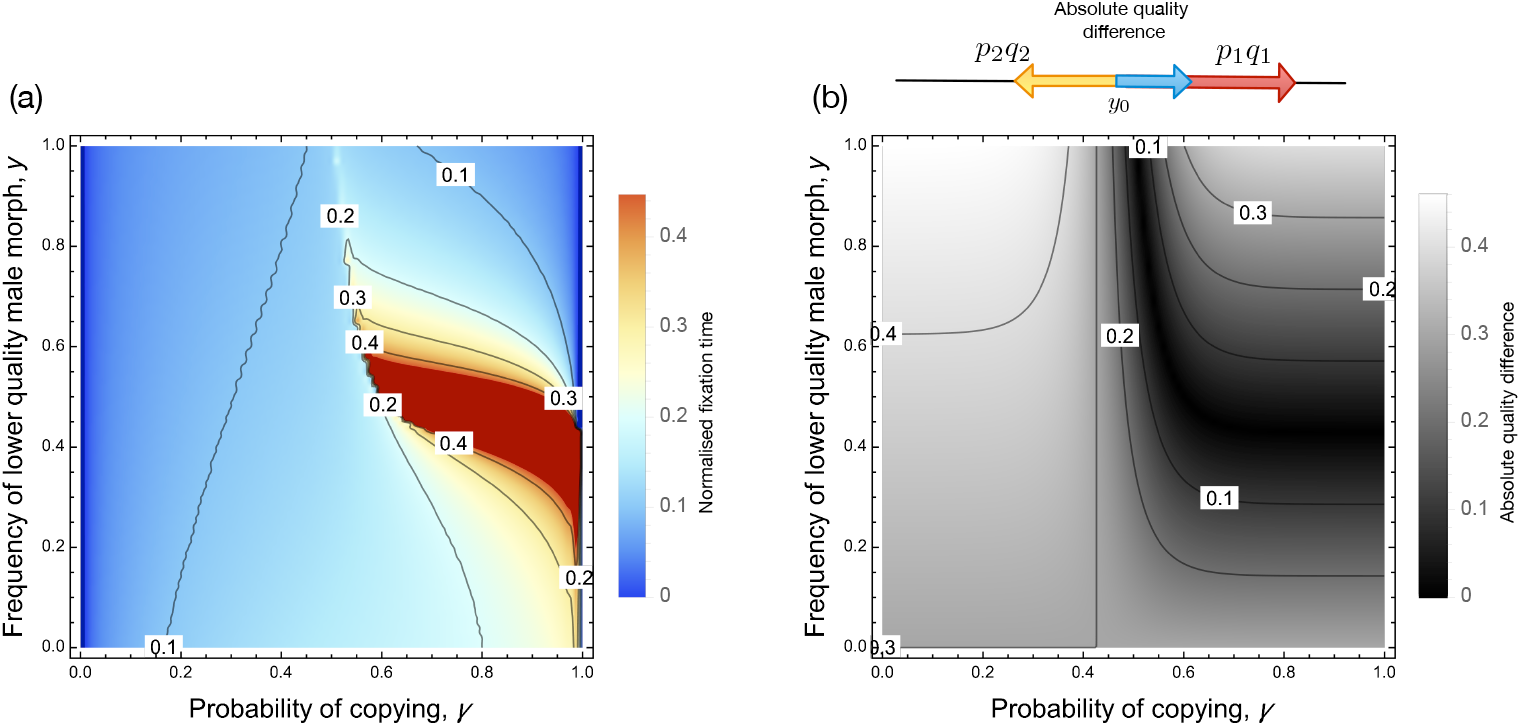
**(a)** Fixation time for the system with two male morphs (Q = [2.0, 2.3], *α* = 0.2, *u* = 0.01, *β* = 20). Comparing to the bifurcation diagram in Figure 2, we observe that the time to fixation is high for regions around the boundary of the basins of attraction. **(b)** Absolute difference between modified quality values. The black band is where the value of the difference is low, and hence the time to reach fixation is high. Parameter values used to compute the numerical solution are given in Table 2. An illustration of how the absolute difference in modified qualities is calculated is given. Magnitudes of the yellow and red vectors are proportional to modified qualities of the two morphs (at the point *y*_0_). Magnitude of the resultant blue vector is the quantity that is plotted.

To visualise the relationship between modified quality differences and fixation, we can imagine vectors that pull the population towards one of the two fixed points (*y* = 0 and *y* = 1). The magnitude of each vector is proportional to the modified quality of a morph (Figure 3b). The difference in the magnitudes of the vectors is proportional to the difference in growth rates, thus indicating the time to reach fixation (Figure 3a). With this exposition of the two morph case we are now able to transition to the relatively more complex three morph case.

### Three Morphs

A system with three male morphs is complex and hard to study experimentally and theoretically. Our model provides valuable insights into the dynamics of the male population with three morphs of different qualities. The results derived from the equations (SI.4) are summarised in Figure 4. The first column shows the phase portrait of the system as the probability of copying (*γ*) changes. When there is no copying (*γ* = 0), there is one stable and two unstable fixed points (at the three vertices), and any initial condition leads to fixation for the morph with the highest quality (morph 3). These fixed points switch stability at higher values of *γ* till all of them are stable. With an increase in copying, new unstable fixed points appear on the boundary and in the interior of the simplex. This allows the population to be monomorphic for each of the three morphs, depending on the initial conditions. The existence of unstable fixed points in the interior and stable fixed points on the vertices implies that any system starting from the interior will move towards one of the vertices with time.

**Figure 4:**
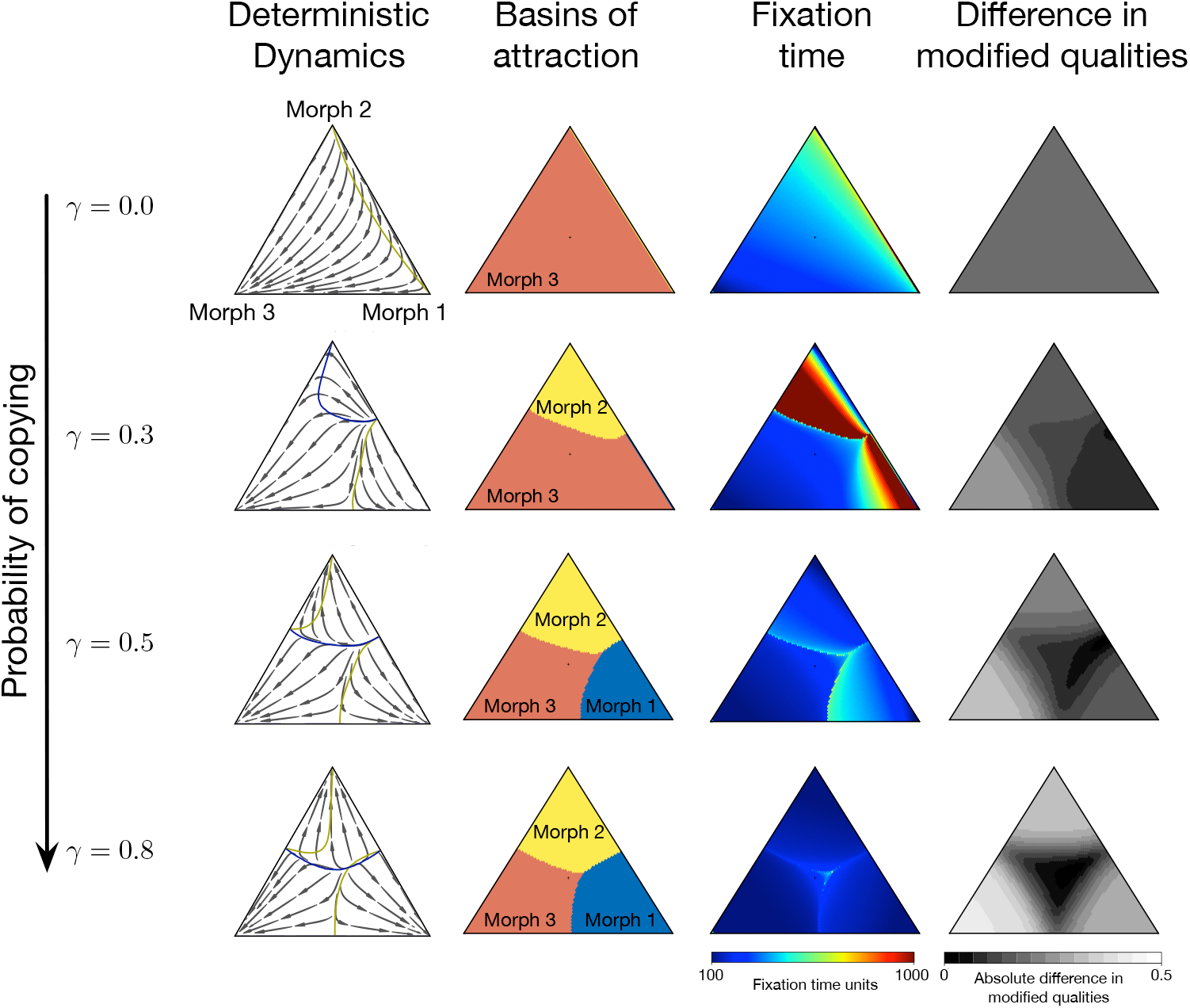
Analysis of the 3-morph system (Q = [2.0, 2.1, 2.3], *α* = 0.2, *u* = 0.01, *β* = 20). Each row corresponds to the indicated value of the probability of copying *γ*. The first column (leftmost) shows the phase portraits of the two-dimensional system. As the value of *γ* increases, we find that new fixed points appear. The second and third columns show the outcomes of the numerical solution of the differential equation for the 3-morph system. We observe that for high values of *γ*, the basins of attraction for the three morphs are almost equal in area. The last column shows the difference in modified qualities, which can be compared to the fixation time plots. The numerical solution is run for 2000 time units.

The second column of Figure 4 shows the basins of attraction for the vertices. With increasing copying probability, the three basins of attraction cover *almost* equal area in the phase space, but even at *γ* = 1, i.e. in the absence of inherent preference, they are not precisely equal. This is because of quality differences and, therefore, differences in growth rates among the subpopulations. Time to fixation (in the third column, Figure 4) is related to the difference in modified qualities (fourth column, Figure 4). The more the difference, the lower is the fixation time. The difference in modified qualities is calculated using vectors, similar to the two-morph case (see Figure SI.1). We take three vectors pointing at each vertex of the morph space, with magnitudes equal to the modified qualities, and compute the resultant magnitude of their sum.

### Search Parameter and Effective Conformism

An infinitely large well-mixed population allows us to make some analytical headway for an arbitrary number *m* of male morphs. The dynamics as described in equations 4 can be analytically solved to give us insights into the qualitative and quantitative nature of the system’s equilibria. We have also performed simulations with the same parameters (such as the probability of copying *γ* and the fraction of matings in the training period *u*).

For the two-morph case, Figure 2 shows the results of these simulations. However, the values of *β* used in the simulation and for obtaining the expected separatrix are different. The reason for this discrepancy lies in the details of the process of mating. In the simulations, as described in Section S.3, the females exhibit inherent or cultural preferences prior to mating. When looking for mates, the females would have to “search” through a certain number of potential mates (by random encounters) before finding the one fitting their preference. This search number in the simulations is a fixed value *s*. The female would give up and settle for the last randomly encountered male after *s* attempts. However, we do not account for this “search parameter” in the deterministic model and analyses. This discrepancy is evident in the value of the copying function *C*.

Mate copying involves copying preferences or switching preferences from inherent ones to learnt ones. Since the search parameter is fixed, all females may not be able to realise their preferences, which leads to lesser switching than expected. We thus replace the copying function *C*(*y*) with an *effective C*(*y*). Consequently, the extent of conformism is reduced. The expression for the effective *C*(*y*) with parameter *s* is as shown below (derivation in Section S.4).

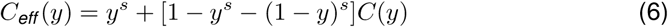

As *s* → ∞ we see that *C*_*eff*_ (*y*) → *C*(*y*). The difference between expected and observed results in Figure 2c is not entirely and quantitatively explained by this relation, which is possibly due to other factors like stochastic effects, limit on the population size, and female population dynamics, which are not included in our numerical model. While this leads us to discuss the benefits and limitations of our model, note that the results are qualitatively the same, and the following inferences still hold.

## Discussion

Our model examines the effect of female mate copying on populations with multiple male morphs, something which has not been explored in previous studies on MC. Results from our model can be used to inform experiments on mating systems with multiple morphs. We demonstrate that a costly or “low quality” morph can go to fixation in the presence of MC. This result is derived for systems with two and three morphs.

Previous theoretical work on MC has looked at dynamics in the presence of two male morphs [28, 16, 30, 29]. In more than one way, our model is highly generalisable. First, it can easily be extended to analyse systems with multiple male morphs. Second, the modified qualities of male morphs (as in Eq. (4)) can be edited to include any other effect on the fitness of males such as ornamentation or intrasexual conflicts [35]. Third, we have shown the effect of different probabilities of copying (*γ*) on the dynamics of the male morphs. This can be used in further studies to show the co-evolution of female copying behaviour and the male population.

We look in detail at the fixation time for the systems under consideration. For the three-morph system, the male population can be represented by a triangular simplex whose vertices represent the monomorphic populations. In the region around the separatrix (the set of unstable fixed points separating the basins of attraction of the vertices), the fixation time is high compared to surrounding regions. Also in this region the difference in modified qualities is very low. This makes intuitive sense because when the difference in growth rates of the morphs is low, it takes longer for the system to change. This has potentially far-reaching consequences. If we include the dynamics of *γ* into the model, then the system could reach a quasi-stable equilibrium under the right circumstances. This would happen when the value of *γ* changes with time such that the fixation time keeps on increasing. We observe non-monotonous behaviour of fixation time for different initial points in the phase space of the three-morph system (Figure 5). The region in the simplex where such behaviour can be observed can account for almost two-thirds of the phase space (the shaded region in Figure 5 right). In this region, the fixation time peaks for a specific values of *γ* (Figure 5 left).

**Figure 5:**
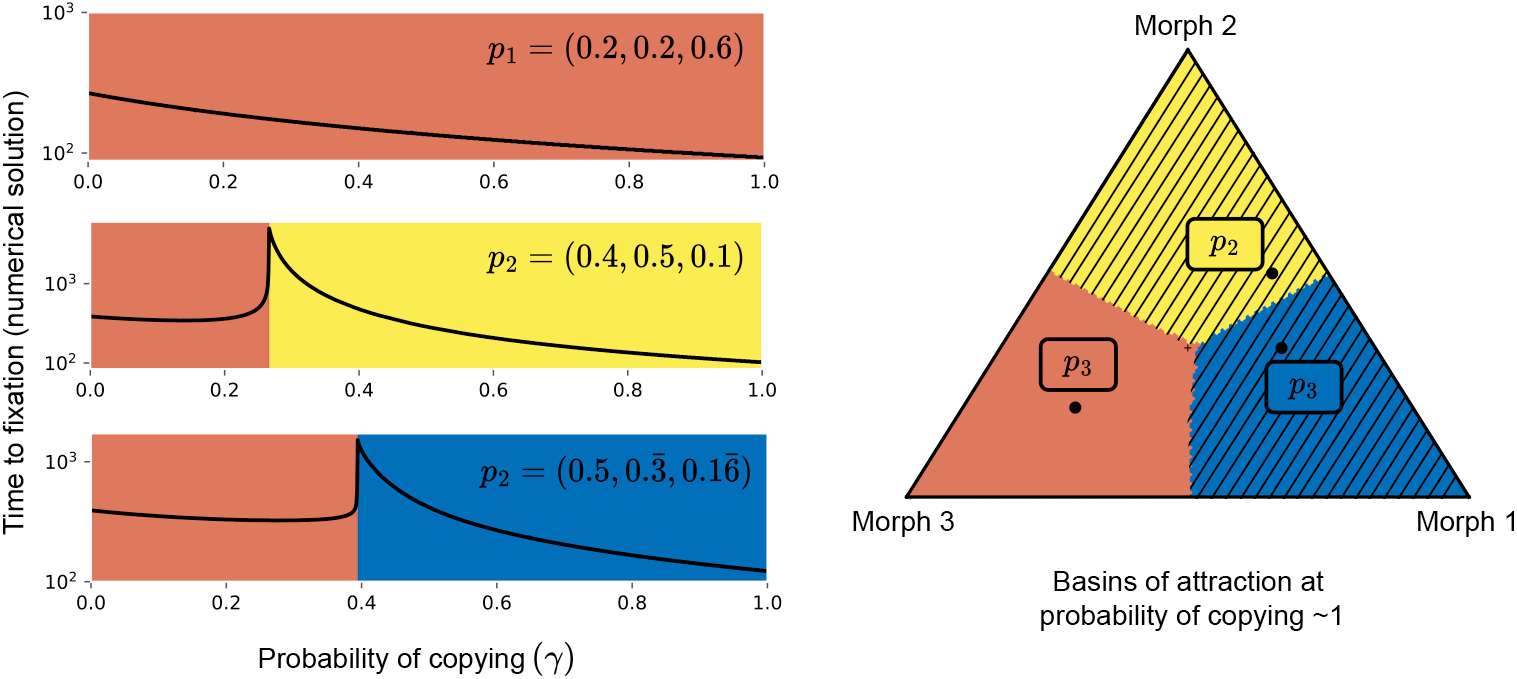
Fixation time for three initial conditions in the phase space as a function of copying probability (Q = [2.0, 2.1, 2.3], *α* = 0.2, *u* = 0.01, *β* = 20). The figure on the right shows the basins of attractions at a probability of copying close to one. For all the points in the shaded region, the time to fixation shoots up (as shown in the plots on the left). This peak occurs when the point switches basins of attraction. This implies that with a changing value of *γ*, the system could reach a quasi-stable polymorphic equilibrium.

In our analysis, we have kept the extent of conformism (*β*) constant. We also noted the difference in *γ* and *β* and how changing them affects conformist behaviour. This can further be extended by varying conformity coefficients over time [34] which can lead to significantly different evolutionary dynamics. Depending on the expected level of conformity, stochastically locally stable equilibria can be observed, which are not possible in the deterministic case, as demonstrated by our model. This postulates a possible mechanism allowing the maintenance of polymorphism in the male population. Connecting this process with multiple morphs as in our study would be a future avenue of research. To incorporate temporal change in the probability of copying, a careful analysis of the costs and benefits of copying behaviour is required. The search parameter is an essential factor for the same. Although all females have to “search” for their preferred male, copying females have an advantage because they are likely to find their preference sooner than non-copying females since they are more likely to switch preference to the most common male in the population. This preference for commonality thus confers a benefit to copying individuals. In this model however, non-copying females have the advantage that they are less likely to mate with low-quality males because it is assumed that inherent preference is in proportion to the quality of male morphs. It is therefore reasonable to assume that the proportion of copying females would reach an equilibrium (see [20]).

We define the function *C*(*y*) as the copying function or the conformism function in our derivations. However, the results remain qualitatively the same for any appropriate function such that the probability of copying preference for the common trait (more precisely, the male morph that mates the most frequently) is more than that of the rarer trait. The “competition” between the preference for commonality and inherent preference is succinctly represented as the difference in modified qualities. Modified qualities account for preference and quality both (Eq. (2)). This equips us with a potent tool for analysing interactions involving mating preferences.

There are many avenues to be explored in modelling mating behaviour like MC, for instance, the effect of spatial distribution of the population where females only copy neighbours. Age, reputation and reliability of model females would have an effect on the frequency of copying [26, 36, 37]. The delay between consecutive observations and the nature of information obtained clearly affect the production of copying behaviour [38, 33, 39]. Future models could consider the effect of inter-generational learning with overlapping generations, and the interplay of multiple observations [34, 38] with multiple morphs. There is still much to explore in mate-copying studies in a co-evolutionary context of multiple male morphs and female preferences and types.

In summary, the generalisability of our model makes it possible for us to analyse systems with multiple male morphs in the context of mate copying and similar conformist behaviours. Using the model, we can edit/add parameters in the expression for modified quality to account for mating preferences, both positive/negative and inherent/socially learnt. Simple conditions for characterising equilibria or quasi-stable coexistence can be derived with modified qualities. Future work can address how the factors mentioned above affect population dynamics and incorporate them into our model.

## Code availability

Appropriate computer code describing the model is available at https://github.com/tecoevo/matecopying_multiplemorphs.

## Acknowledgements

CSG acknowledges the visiting fellowship from IAST that helped start this project. SN acknowledges IAST funding from the French National Research Agency (ANR) under the Investments for the Future (Investissements d’Avenir) program (ANR-17-EUR-0010). Funding from the Max Planck Society is graciously acknowledged.

## Supplementary material

### S.1 Fixed points in the two-morph system

Following our model (Eq (4)), the system of dynamical equations for a population with two morphs is:

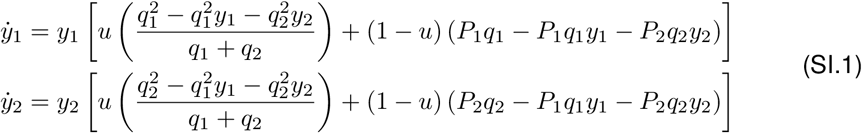

where, *y*_*i*_ and *q*_*i*_ are the subpopulation frequencies and quality values for the two morphs (*i* = 1, 2) respectively. The preference factors for the morphs are *P*_*i*_ = (1 − *γ*)*q*_*i*_*/*(*q*_1_ + *q*_2_) + *γαC*(*y*_*i*_). The copying function *C*(*y*_*i*_) is as defined in equation (3). The training period (as described in the section Training Period), is a fraction *u* of the total number of matings in a generation.

The state space is one-dimensional, and we look at one of the two equations above - the one corresponding to the morph with lower quality, let it be *y*. The parameter of interest to look at is *γ* (copying probability), and we study the behaviour of the system for a range of *γ*. Other parameters are fixed at values given in Table 2. For our analysis, we take the quality values to be close to each other. For larger differences in quality, inherent preference is expected to dominate, resulting in the fixation of the higher quality morph. Additionally, a large difference in quality implies a large difference in number of offspring of the two morphs. This would contribute to the higher quality morph subpopulation going to fixation. We can find the threshold quality difference below which the higher quality morph subpopulation can go *extinct*. Since the sum of 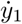 and 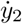 is 0 (constant population), it is clear that both *y* = 0 and *y* = 1 are fixed points that exist for all values of the parameter *γ*. To find other fixed points, we find the values of *y* at which equation (SI.1) equals 0, i.e.

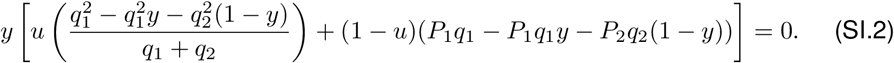

Solving this gives us equation (5), the expression for the internal fixed point, where

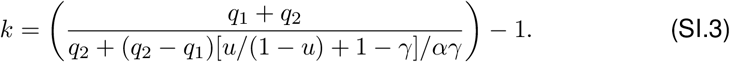

Whenever the value of *k* is non-negative, the internal fixed point *y*^∗^ exists. Its value depends on the parameter *γ* (as shown in Figure 2).

#### Linear Stability Analysis

Let us call equation (SI.2) *f*(*y*). We find that *f*^′^(0) *<* 0 for all values of *γ* and fixed values for other parameters (as mentioned in Figure 2). The stability of the other fixed point(s), *y* = 1 and *y* = *y*^∗^, depends on the sign of *f* ^′^ at these points. We obtain that *f*^′^(*y*^∗^) *>* 0 for all “valid” values of *y*^∗^ i.e., *y* between 0 and 1 that satisfy equation (5).

At the birth of *y*^∗^, there exists a bifurcation in the *γ* − *y* phase space (illustrated in Figure 2). At this bifurcation point, *f* ^′^(1) changes its sign from positive to negative and becomes a stable, attracting fixed point, which means that there is a region in the phase space from where the lower quality morph can go to fixation. Code to check the derivative of *f*^′^ at different points is provided.

**Figure SI.1:**
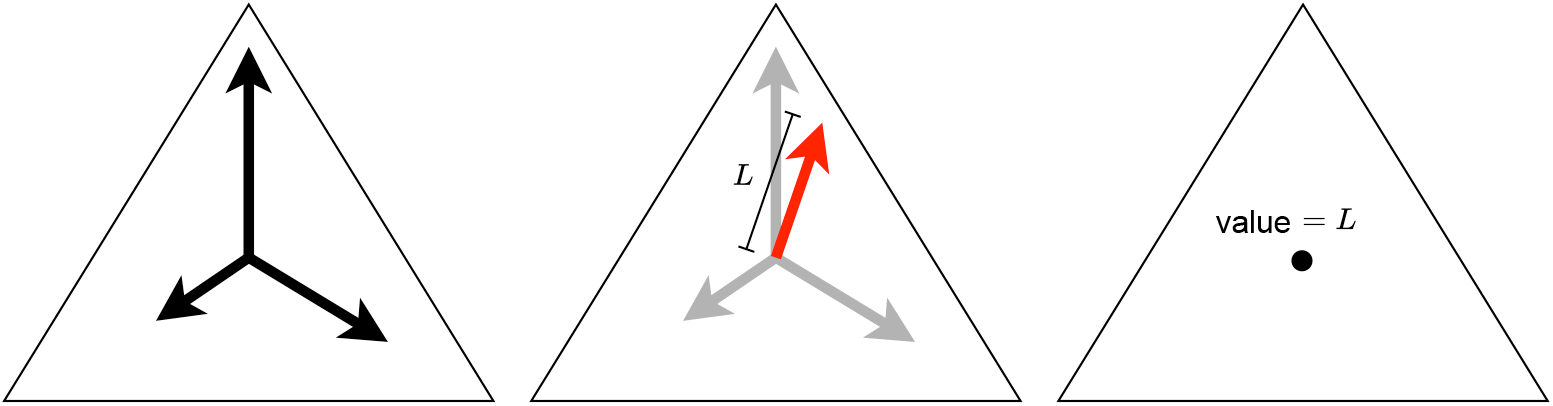
An illustration of how the magnitude of difference in modified qualities is computed. The black vectors are the modified qualities corresponding to each morph in the system. The red vector is the resultant of the three and the value of the difference at the point shown is equal to the magnitude of the resultant vector.

### S.2 Fixed points in the three-morph system

Analogous to the two morphs, the system of dynamical equations for a population with three morphs is:

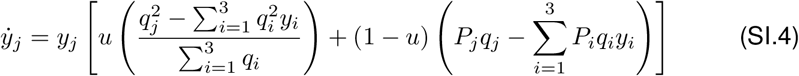

All the parameters and symbols are as defined before for the two-morph case but now for morphs *j* = 1, 2, 3. The phase space is two-dimensional, so we can pick 2 out of the 3 variables in the equations above. We will look at the lowest and highest quality morph populations, (*y*_1_, *y*_3_). The three vertices of the simplex, (0, 1), (1, 0), (0, 0) are fixed points. Their stability varies as the value of *γ* changes. For example, at *γ* = 0, (0, 1) and (0, 0) are both stable fixed points while (1, 0) is unstable fixed point. Other fixed points are born for higher values of *γ* (see Figure 4).

### S.3 Simulation

The behaviour of our system (with *m* morphs) in the limit of an infinite, well-mixed population is estimated by equation (4). We use a specific functional form of the preference factor, which multiplicatively enhances or diminishes fitness is hypothesised to be as given in Eq (2). To support the results of our model and understand the consequences of the assumptions made, we run a simulation of the system with the parameters listed in Table 2 with a finite population of 200 individuals. The results are averaged over a 100 independent runs.

#### One Run

One run of the simulation involves 100 generations. The first generation starts with an initial frequencies of the *m* male morphs. At the end of each generation, the output is a new vector of male morph frequencies. Copying in the next generation is then according to the switching probabilities obtained using these new frequencies. The initial frequency of females that prefer a particular male morph is proportional to the quality of the morph. The higher the quality of a morph, the higher is the number of females that inherently prefer it. The distribution of female preferences changes within a generation but not across generations, because we have assumed that learnt preferences are not heritable.

#### One Generation

One generation of the simulation involves 100 matings, which is equal to the number of females in the population. The first *n* matings constitute the training period, during which all females collect information - specifically the ratio of total matings that involve each male morph. There is no mate copying and thus females preference is inherent and frequency independent.

At the beginning of each generation, females prefer males in proportion to their fitness. For all the matings to follow after the training period, females choose to copy, with a probability *γ*.

#### One Mating

For each mating in the copying period, a female is chosen at random. If this female chooses to copy, it can switch its preference according to the switching probabilities calculated using the information collected during the training period (Equation (3)).

The female then encounters mates at random. It can either accept or reject the encountered male depending on its preference. This process of “searching” for the preferred morph occurs *s* number of times. This means that the female will give up and accept the *s*^th^ male as its mate (if it does not find its preferred mate before).

Since the population is fixed, the number of deaths is equal to the number of births, which is determined by the quality of the male morph. If *q*_*i*_ is the quality of the chosen male, then the number of offspring is sampled from an alpha distribution with mean *q*_*i*_. Deaths occur at random.

#### Caveats

1. Social learning occurs within a generation. The population is well-mixed, so all females copy using the same information.
2. Females gather social information during the training period only.
3. Learnt preferences are not inherited, and inherent preferences are in proportion to male morph qualities. This implies that female preferences at the start of each generation are the same.
4. Female births and deaths occur such that it does not affect the frequency of females that prefer a male morph. This assumption works if the difference in male qualities is small because births and deaths both occur in proportion to male morph qualities.
5. The number of offspring from a mating is at least one. Therefore, male morph quality cannot be less than one.
6. Switching probabilities are independent of female preference prior to copying. This is not true for one mating, but we show that it works on an average. If *f*_1_ is the frequency of females that prefer morph 1, then this frequency after a generation of copying will be

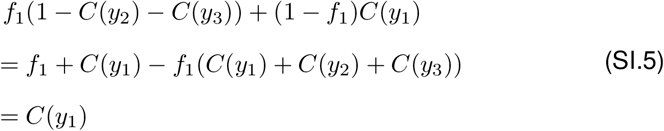
7. The value of the search parameter *s* is fixed. In our simulation, both copying and non-copying females meet at most *s* males before choosing their mate, but in general the value of *s* could be different for these two types of females. The fixed value is also finite, which means that all females may not mate with their preferred male.

### S.4 Search Parameter and Effective Conformism Derivation

The function *C*(*y*) encodes the extent of conformism. Due to the presence of a fixed search parameter, the extent of conformism is effectively reduced. Equation (6) gives an expression for the *effective* extent of conformism. For the two-morph case, we consider the following conditional probabilities (here, 1 and 2 referring to the male morph labels and *y*_*i*_ to their respective frequencies),

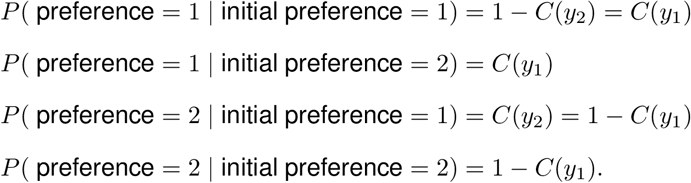

Let *y* = *y*_1_, since all the required expressions are in terms of *y*_1_. If we assume that every female gets its preference, that is it keeps looking till it finds it preference, then,

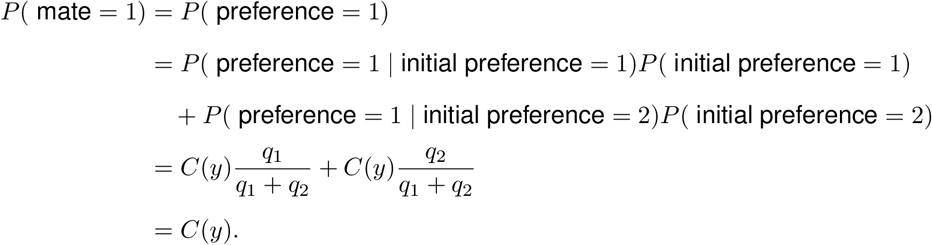

**Figure SI.2:**
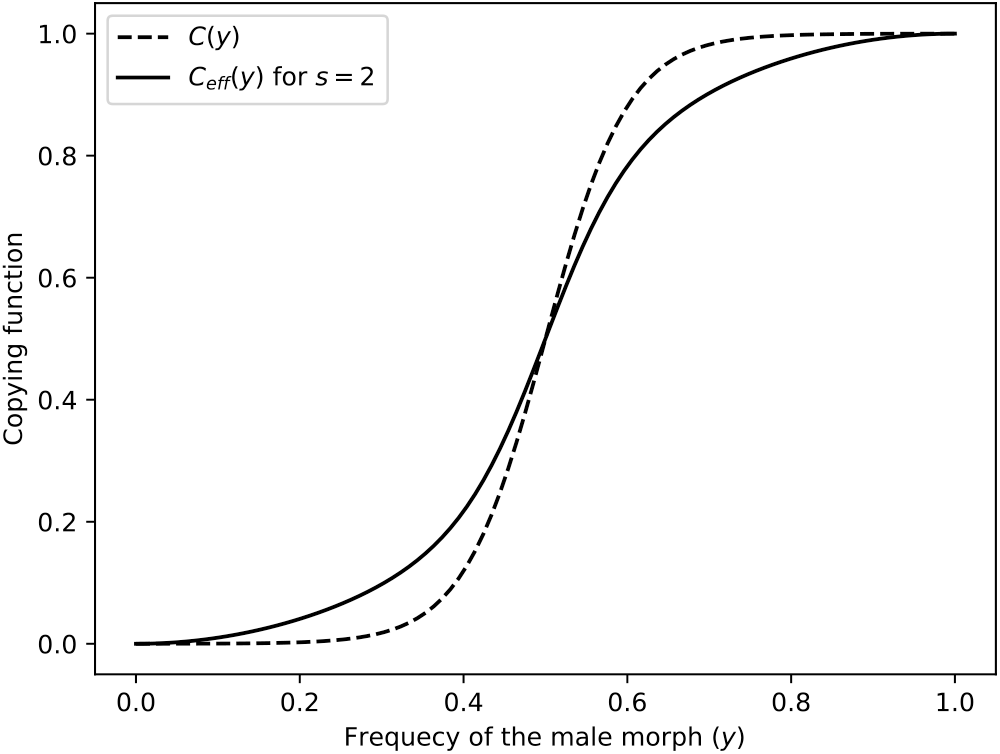
Effective copying function *C*_eff_(*y*) for search parameter *s* = 2. The dashed line shows the copying function without adjusting for the search parameter. Note that for the same value of *β*, effective copying function shows a higher extent of conformism than *C*(*y*).

Effectively, the probability of mating with the preferred morph is reduced if the number of times a female can “search” is fixed at a finite value *s*.

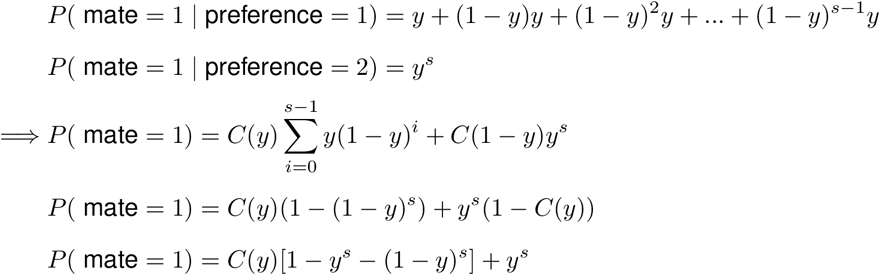

Therefore, the frequency of mating with morph 1 is less than the frequency of females preferring morph 1. This reduction is apparent in Figure SI.2

